# Optimization of denoising and filtering parameters of DADA2 for QIIME2 amplicon metagenomics data analysis

**DOI:** 10.1101/2025.06.24.661437

**Authors:** Moirangthem Goutam Singh, Romi Wahengbam

## Abstract

**Background:** High-throughput sequencing generates vast data, often containing low-quality bases, chimeras, and artifacts that can mislead taxonomic classification and diversity assessments. DADA2 enhances taxonomic resolution by excluding low-quality bases and optimizing ASV inference. Proper truncation reduces computational load while maintaining key hypervariable regions for accurate classification. In this study, we examine the effect of various truncation lengths during the DADA2 analysis in ensuring statistical robustness and improving the reliability of microbial community profiling in ecological and environmental studies.

**Results:** Truncation of read length improves the quality reads recovery rate, and preserves microbial diversity in the V4 hypervariable region of Illumina paired-end reads.

**Conclusion:** Incorporating the best truncation length strategy optimizes the reads recovery and preserves the richness and evenness of microbial communities.

## Background

In microbial ecology and environmental biology, analyzing microbial communities through high-throughput sequencing has revolutionized our understanding of biodiversity, ecological interactions, and ecosystem functioning. QIIME 2 (Quantitative insights into microbial ecology), one of the most widely used platforms for analyzing and interpreting microbiome data, offers comprehensive tools for processing and visualizing large sets of sequencing data. Among the various preprocessing steps, trimming the sequence length of reads is crucial to ensure the accuracy and reliability of downstream analyses [1, 2]. High-throughput sequencing technologies, such as Illumina sequencing, generate vast amounts of sequencing data that provide insights into the composition and function of microbial communities. However, raw sequences often come with inherent issues such as low-quality bases, chimeras, and non-target (host or contaminant) sequences. These factors can dramatically affect the quality of the data and, consequently, the interpretations drawn from it [3]. Quality control is a fundamental aspect of sequence analysis that determines the integrity and validity of results. Trimmed sequences help polish the dataset’s quality, allowing for better detection of true microbial diversity and accurate taxonomic assignment [4]. Low-quality sequences can introduce noise into the dataset, potentially leading researchers to erroneous conclusions regarding the composition and structure of microbial communities. By trimming sequences, researchers can remove parts of reads that do not meet quality standards, significantly increasing the overall data quality [5]. Sequencing artifacts, including errors introduced during the sequencing process, can result in misinterpretations of microbial community structure. This can involve substitutions, insertions, deletions, or biases associated with specific sequences [6]. Trimming identified low-quality bases from the beginning and end of sequences helps eliminate potential mismatches during alignment and reduces the likelihood of assigning incorrect taxonomy to operational taxonomic units (OTUs) or amplicon sequence variants (ASVs) [7]. Effective trimming can also reduce the creation of chimeric sequences, which arise when two sequences join together, leading to false inflation of biodiversity estimates [8].

Performance in downstream statistical analyses relies heavily on the quality of input data. Tools used for diversity analyses, such as alpha and beta diversity metrics, depend on accurately representing community richness and evenness. If sequences are left untrimmed, analyses can produce misleading results, with potential increases in false positives and negatives [9]. For instance, alpha diversity measures such as Shannon or Simpson indices can be disproportionately influenced if low-quality sequences are included in these calculations. Addressing these concerns through effective trimming ensures that all statistical assumptions and models are based on high-quality, reliable data. The specifics of trimming often involve determining an appropriate “cut-off” length for sequences. This cut-off is typically chosen based on several criteria. The quality score associated with each base in a sequence is a critical determinant in the decision to trim. The Phred score, commonly used in sequencing data, provides a logarithmic representation of base call accuracy. Sequences that fall below a predetermined threshold (for example, Q20 or Q30) are often candidates for trimming or exclusion [10]. Sequences that are shorter than a defined minimum length after trimming may not contain enough information for accurate taxonomic assignment. Trimming sequences to a consistent length ensures that subsequent analyses within a study are based on comparable data breadth across samples [2]. For paired-end sequencing, overlapping regions between forward and reverse reads may need to be trimmed to yield optimal merged reads. This step enhances the accuracy of read merging and helps reduce the generation of uninformative sequences [8]. In many studies, particular regions (like the 16S rRNA gene) are targeted for amplification. Trimming ensures that only the regions of interest are retained, mitigating the influence of extraneous information that can arise from untrimmed reads [3].

Within the QIIME 2 framework, DADA2 plays a crucial role in the analysis pipeline, facilitating accurate taxonomic classification and community structure analysis. The integration of DADA2 in QIIME enables researchers to obtain more reliable and reproducible results in ecological studies of microbial populations [2]. Truncating sequences to a specific length during DADA2 analysis in QIIME 2 is an essential strategy for improving the quality and accuracy of microbial community profiling. This process involves shortening the sequences, particularly the ends where low-quality bases may occur, which can lead to sequencing errors and affect downstream analyses. The effectiveness of this technique is underscored by several key factors that support its application in microbiome research.

One of the primary benefits of truncation is the significant reduction of sequencing errors. High-throughput sequencing technologies, while powerful, often generate low-quality bases at the ends of the read, especially in the context of longer reads. These base call inaccuracies can mislead taxonomic assignments and inflate measures of diversity. DADA2 improves its performance by using quality scores to inform truncation length decisions [2]. By defining a truncation length that excludes low-quality bases, researchers ensure that only the most reliable data are retained for analysis.

DADA2 employs a denoising algorithm that is sensitive to the quality of the input sequences. When poor-quality bases are present, the algorithm may struggle to accurately distinguish true sequences from artifacts or noise [7]. Truncating the sequences to a specific length helps create a more stable input by excluding these problematic ends, thereby allowing DADA2 to generate more reliable amplicon sequence variants (ASVs) representative of the actual microbial community. It has been demonstrated that appropriate truncation, in conjunction with quality filtering, significantly enhances the accuracy of ASV inference [1].

Truncating sequences to a length that maintains the core hypervariable regions (e.g., V3-V4 or V4) enables effective taxonomic resolution while reducing the likelihood of including less informative or error-prone sequence data. Many studies have shown that retaining specific length ranges where the sequence variability is high contributes positively to the precision of taxonomic classification across complex microbial communities [10]. This approach is particularly useful for distinguishing closely related taxa, allowing for more robust interpretations of microbial diversity and community structure.

Truncating sequences also streamlines the computational burden in downstream analyses. Longer reads can lead to increased processing times and memory usage. By specifying an optimal truncation length, researchers can reduce the complexity of the data, making analyses such as diversity assessments or statistical comparisons more manageable [11]. This efficiency is especially important during large-scale microbiome studies, where datasets can become quite extensive.

This study aims to contribute to the body of knowledge by conducting a comprehensive analysis of truncation length through a comparative lens. We will identify best practices by assessing already published datasets and provide evidence-based recommendations for determining optimal truncation lengths. The discussion will highlight how these practices enhance the clarity and precision of research findings and ensure that valuable information is not inadvertently discarded.

## Results

### Quality control Assessment

A rigorous quality assessment of the sequences was conducted before the execution of DADA2 and subsequent analyses (Fig. 1). Forward reads exhibited high-quality sequence bases at the onset of the sequencing cycle, with a notable decline in quality towards the end (Fig. 1a). In contrast, reverse reads displayed comparatively lower base quality throughout the cycle (Fig. 1b). To evaluate the impact of trimming, a series of truncation lengths—specifically 0, 170, 175, 180, 185, 190, 195, and 200 bases—were applied to both forward and reverse reads.

**Fig. 1.**
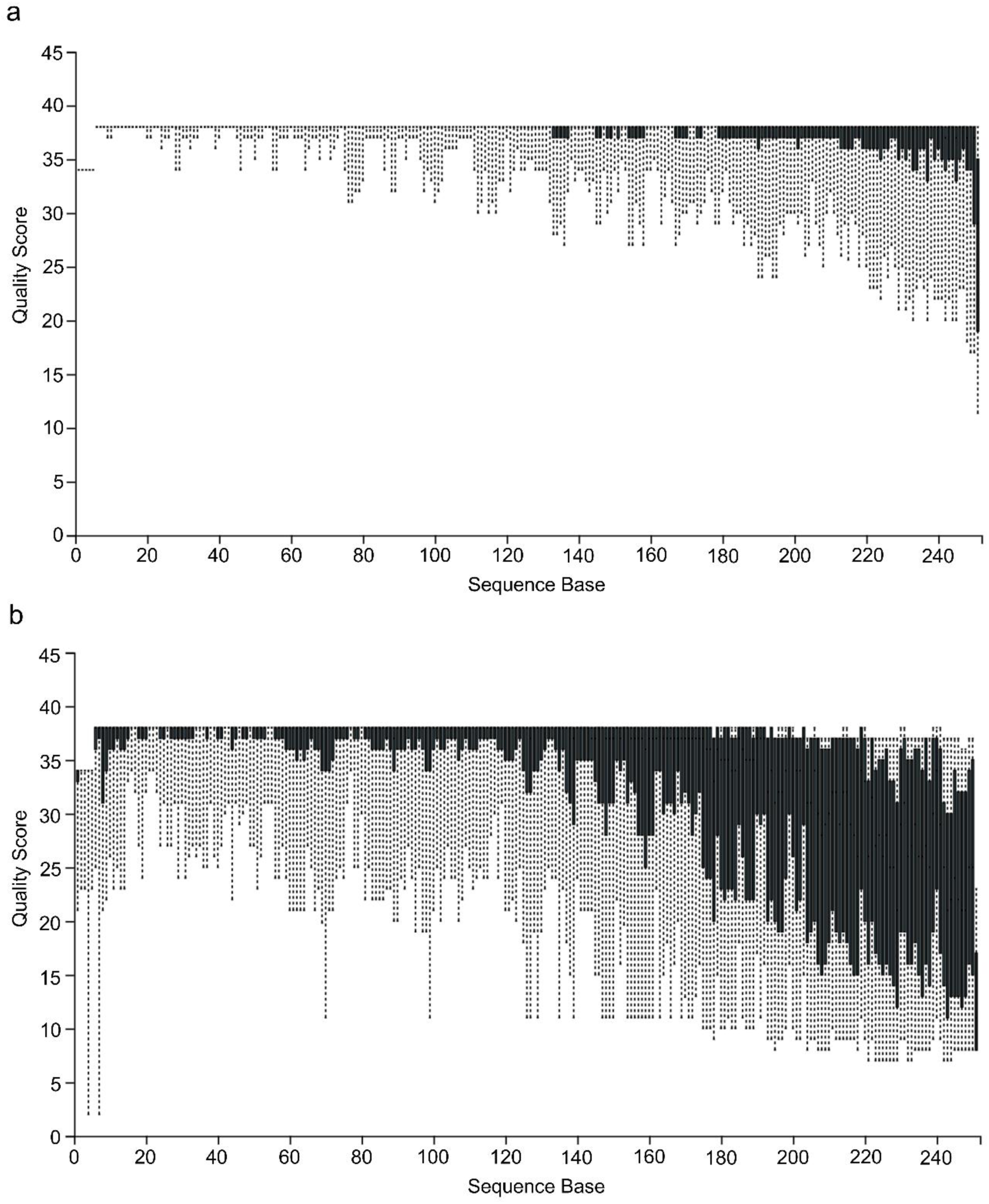
Representative box plot of quality scores over positions in sequenced reads generated using a random sampling of 10000 out of 2277091 sequences without replacement. (**a**) quality distribution for forward reads (**b)** quality distribution for reverse reads. The upper whisker, top of box, middle box, bottom box, and lower whisker represent 91^st^, 75^th^, 50^th^, 25^th^, and 9^th^ percentile quality scores, respectively.

### Good reads count

The DADA2 analysis unequivocally indicated that the parameter with no truncation (trunc_length_0) produced the highest counts of filtered reads, denoised reads, merged reads, and non-chimeric reads (Tab. 1). Subsequent truncation lengths (170, 175, 180, 185, 190) resulted in progressively reduced read counts, with trunc_len_200 yielding the lowest. The percentage difference between input and filtered reads was remarkably minimal for trunc_length_0 at 25.18%, while approximately 45% of denoised reads were lost during the merging of paired-end sequences (Fig. 2). Importantly, trunc_len_175 and trunc_length_170 accounted for losses of 60.31% and 59.33% of input reads, respectively, during the filtering phase. Therefore, the no-truncation parameter decisively yielded the maximum counts of non-chimeric reads

**Table 1.**
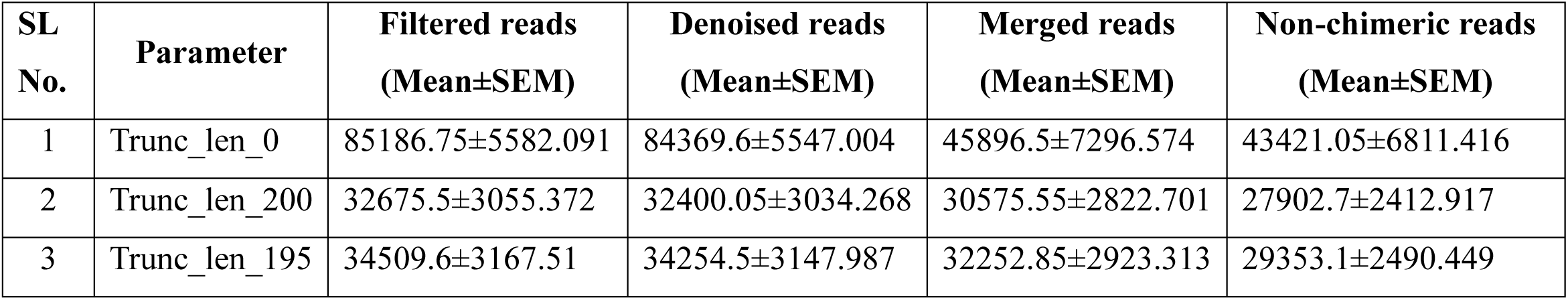
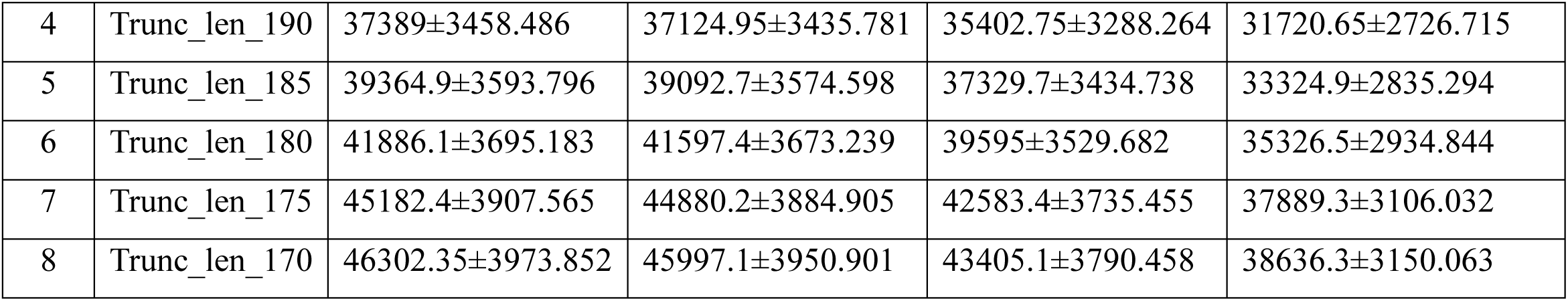
Filtered statistics of reads after DADA2 filtering.

**Fig. 2.**
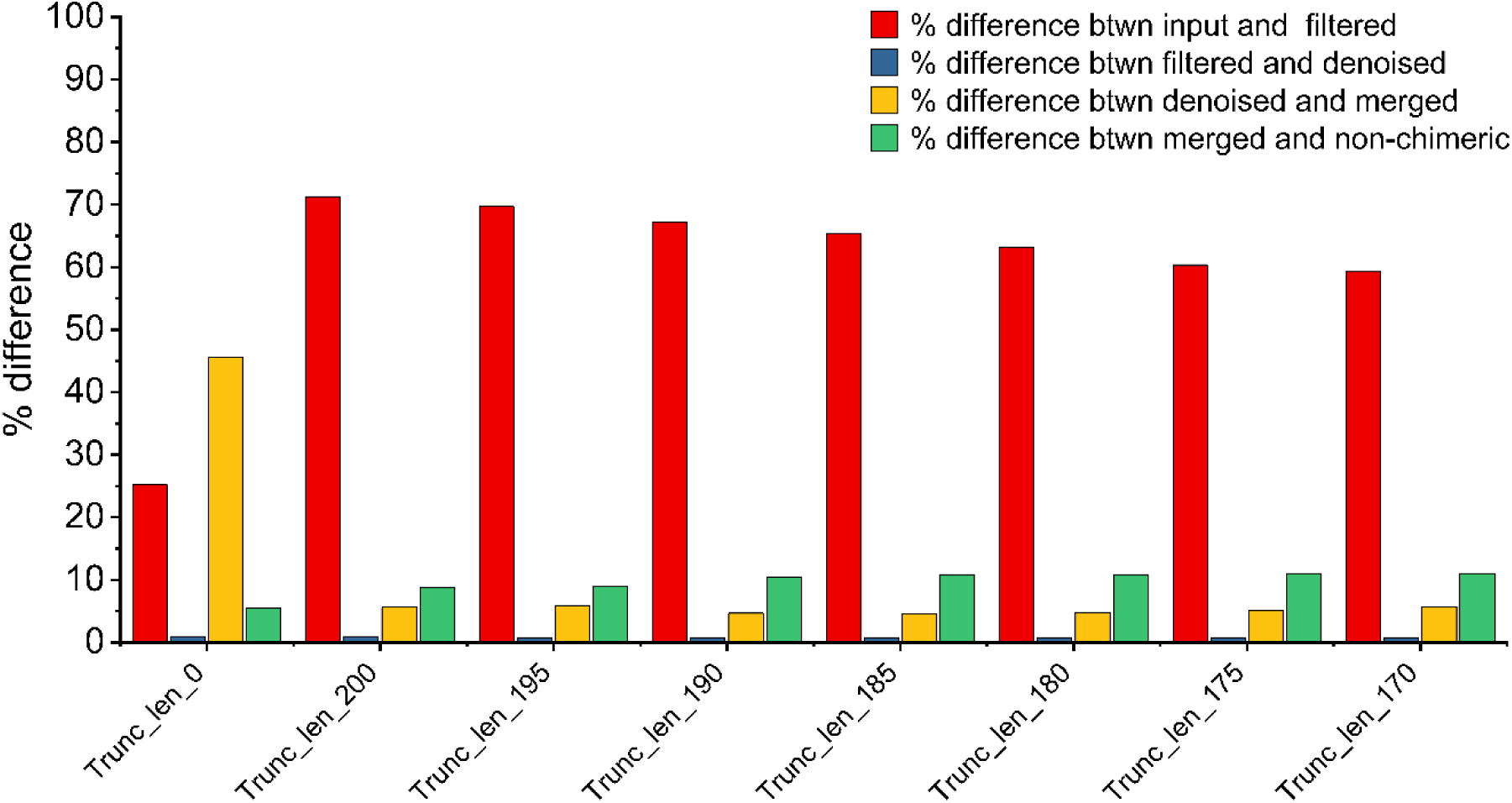
Percentage difference between different filtering step in DADA2 denoising.

### OTU clustering, taxonomic classification, removal of singleton, and doubleton, and filtering of low-frequency and low-abundant features and frequencies

The quantification of frequencies and features was methodically assessed at every stage of OTU clustering, taxonomic classification, removal of singletons and doubletons, and filtering of low-frequency and low-abundant features. Trunc_len_0 emerged with the highest number of frequencies, followed closely by trunc_len_170 and trunc_len_175, while trunc_len_200, as expected, recorded the lowest (Tab. 2). Furthermore, the maximum number of features was definitively observed at trunc_len_195, featuring 359 identifiable features (Tab. 3).

**Table. 2:** Number of feature frequencies after various filtering step viz: DADA2 denoising, OUT Clustering, Taxonomic classification, singleton and doubleton filtering, low frequency filtering and low abundant filtering.

| SL No. | Parameter | DADA2 denoising | OUT Clustering | Taxonomic classification | Singleton and doubleton filtering | Low frequency filtering | Low abundant filtering |
| --- | --- | --- | --- | --- | --- | --- | --- |
| 1 | Trunc_len_0 | 868421 | 868421 | 867971 | 867964 | 862495 | 862495 |
| 2 | Trunc_len_200 | 558054 | 558054 | 557860 | 557852 | 552654 | 552654 |
| 3 | Trunc_len_195 | 587062 | 587062 | 586821 | 586815 | 581329 | 581329 |
| 4 | Trunc_len_190 | 634413 | 634413 | 634151 | 634147 | 628107 | 628107 |
| 5 | Trunc_len_185 | 666498 | 666498 | 666230 | 666228 | 659789 | 659789 |
| 6 | Trunc_len_180 | 706530 | 706530 | 706246 | 706246 | 699535 | 699535 |
| 7 | Trunc_len_175 | 757786 | 757786 | 757533 | 757531 | 750570 | 750570 |
| 8 | Trunc_len_170 | 772726 | 772726 | 772466 | 772461 | 765459 | 765459 |

**Table. 3:** Number of features after different filtering steps viz: DADA2 denoising, OUT Clustering, Taxonomic classification, singleton and doubleton filtering, low-frequency filtering, and low abundant filtering.

| SL No. | Parameter | DADA2 denoising | OUT Clustering | Taxonomic classification | Singleton and doubleton filtering | Low frequency filtering | Low abundant filtering |
| --- | --- | --- | --- | --- | --- | --- | --- |
| 1 | Trunc_len_0 | 460 | 405 | 391 | 386 | 219 | 219 |
| 2 | Trunc_len_200 | 718 | 614 | 607 | 603 | 359 | 359 |
| 3 | Trunc_len_195 | 716 | 614 | 606 | 603 | 358 | 358 |
| 4 | Trunc_len_190 | 726 | 623 | 614 | 612 | 354 | 354 |
| 5 | Trunc_len_185 | 732 | 624 | 615 | 614 | 352 | 352 |
| 6 | Trunc_len_180 | 732 | 624 | 615 | 613 | 350 | 350 |
| 7 | Trunc_len_175 | 722 | 609 | 601 | 600 | 347 | 347 |
| 8 | Trunc_len_170 | 719 | 609 | 601 | 598 | 346 | 346 |

### Diversity analysis

Alpha diversity analysis is a pivotal component in microbiome research, offering insights into the richness and evenness of microbial communities within individual samples. High diversity is often associated with ecosystem stability, resilience, and functionality, particularly in gut microbiota studies, whereas low diversity has been linked to dysbiosis and various diseases, including inflammatory bowel disease (IBD), obesity, and diabetes. In this study, while trunc_len_0 yielded the highest number of good reads, it correspondingly revealed the lowest alpha diversity (Fig. 3a), as also indicated by the rarefaction curve presented in Fig. 4a. Alpha diversity was observed to be greater in group 1 than in group 2, a finding that diverges from conventional expectations. The remaining truncation parameters displayed comparable alpha diversity indices, and the rarefaction curves illustrated a consistent pattern in the relationship between richness and sequencing depth (Fig. 3 and 4).

**Fig. 3.**
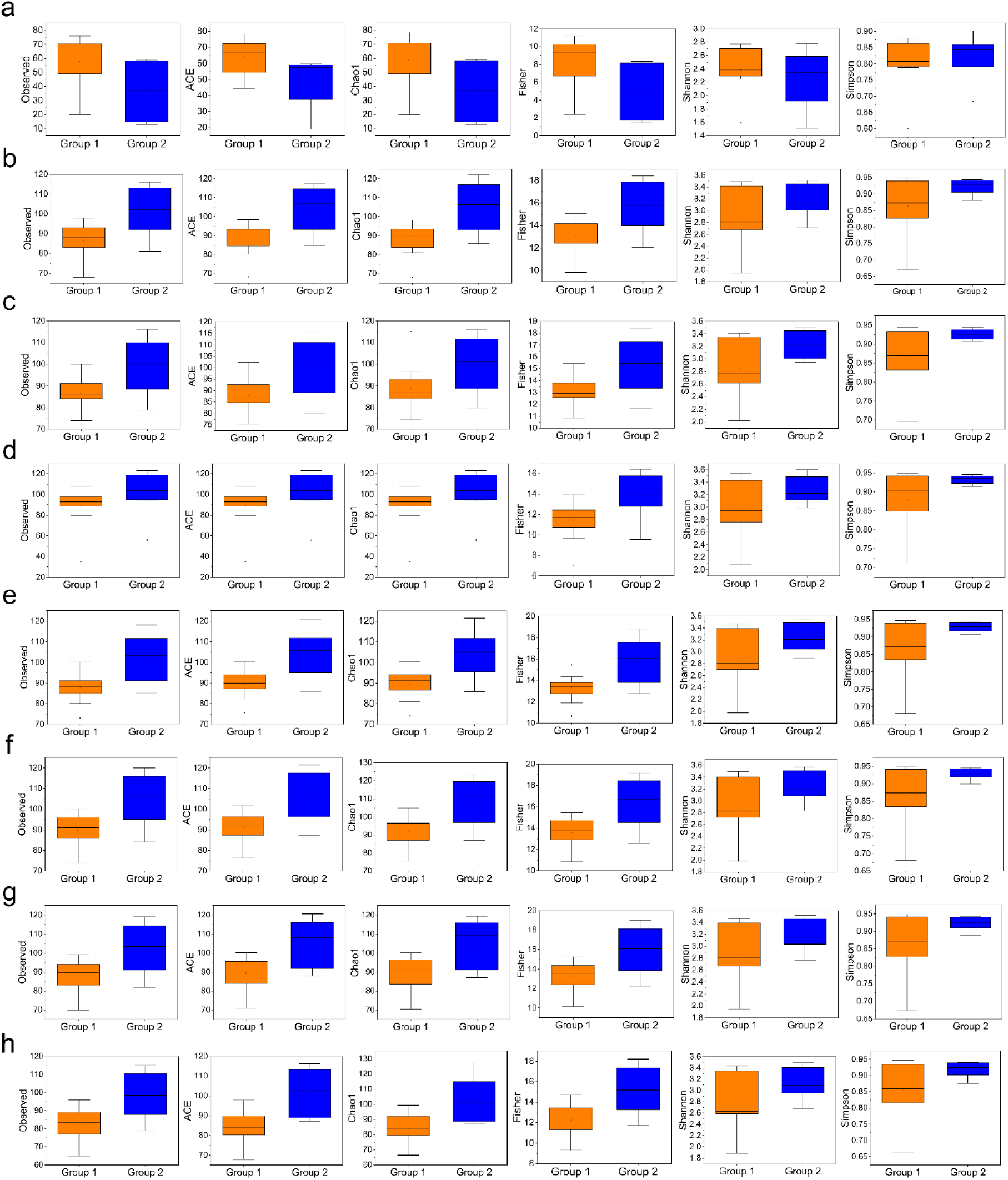
Distribution of alpha diversity indices of different truncation length during DADA2 analysis (**a**) truncation length at 0 (**b**) truncation length at 200 (**c**) truncation length at 195 (**d**) truncation length at 190 (**e**) truncation length at 185 (**f**) truncation length at 180 (**g**) truncation length at 175 (**h**) truncation length at 170.

**Fig. 4.**
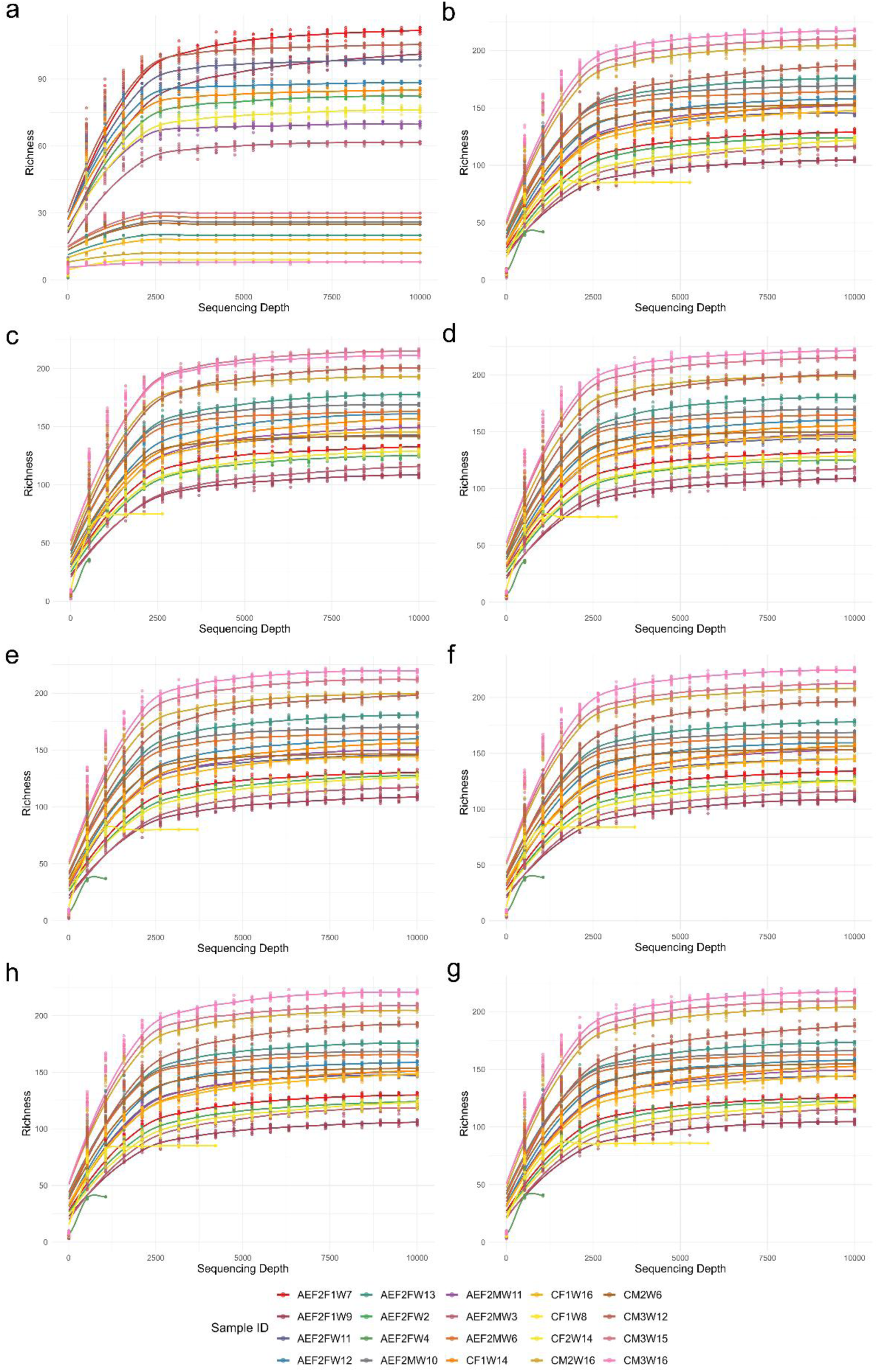
Alpha rarefaction curves (**a**) truncation length at 0 (**b**) truncation length at 200 (**c**) truncation length at 195 (**d**) truncation length at 190 (**e**) truncation length at 185 (**f**) truncation length at 180 (**g**) truncation length at 175 (**h**) truncation length at 170.

### Taxa abundance

Relative abundance at the phyla level revealed three distinct and significant patterns across the various truncation parameters. In the absence of truncation (trunc_len_0), the phylum Firmicutes_D was prominently dominant, followed by Bacteroidota and Firmicutes_A. Conversely, for trunc_len_200, trunc_len_195, trunc_len_190, trunc_len_185, and trunc_len_180, Firmicutes_A was the predominant phylum, succeeded by Firmicutes_D and Bacteroidota. Remarkably, in trunc_len_175 and 170, Bacteroidota emerged as the more prominent phylum compared to Firmicutes_D (Fig. 5). ANCOM comparisons conclusively revealed significant abundance of Bacteroidota and Firmicutes_D in trunc_len_0, whereas Bacteroidota and Firmicutes_B were significantly more prevalent in trunc_len_200, 185, 180, 175, and 170 for group 2 (Fig. 6 a, b, e, f, g, h). Conversely, the phyla Proteobacteria and Verrucomicrobiota showed significant reductions across all truncation parameters except for trunc_len_0 (Fig. 6). This disparity at the phylum level was further substantiated by the analysis of the top 20 species. The highest species-level relative abundances were definitively documented in trunc_len_0, followed by trunc_len_170, 200, 175, 180, 185, 195, and 190. The increased relative abundance in trunc_len_0 predominantly stemmed from the significant presence of species such as *Phocaeicola sartorii*, *Cryptobacteroidetes* sp.009774765, and CAG-485 sp002493045 from the phylum Bacteroidota, as well as *Limosilactobacillus* spp. and *Faecalibaculum rodentium* from the phylum Firmicutes_D (Fig. 7). This underscores the critical influence of sequencing parameters on microbial community structure and diversity, emphasizing the need for careful consideration of truncation strategies in microbiome studies.

**Fig. 5.**
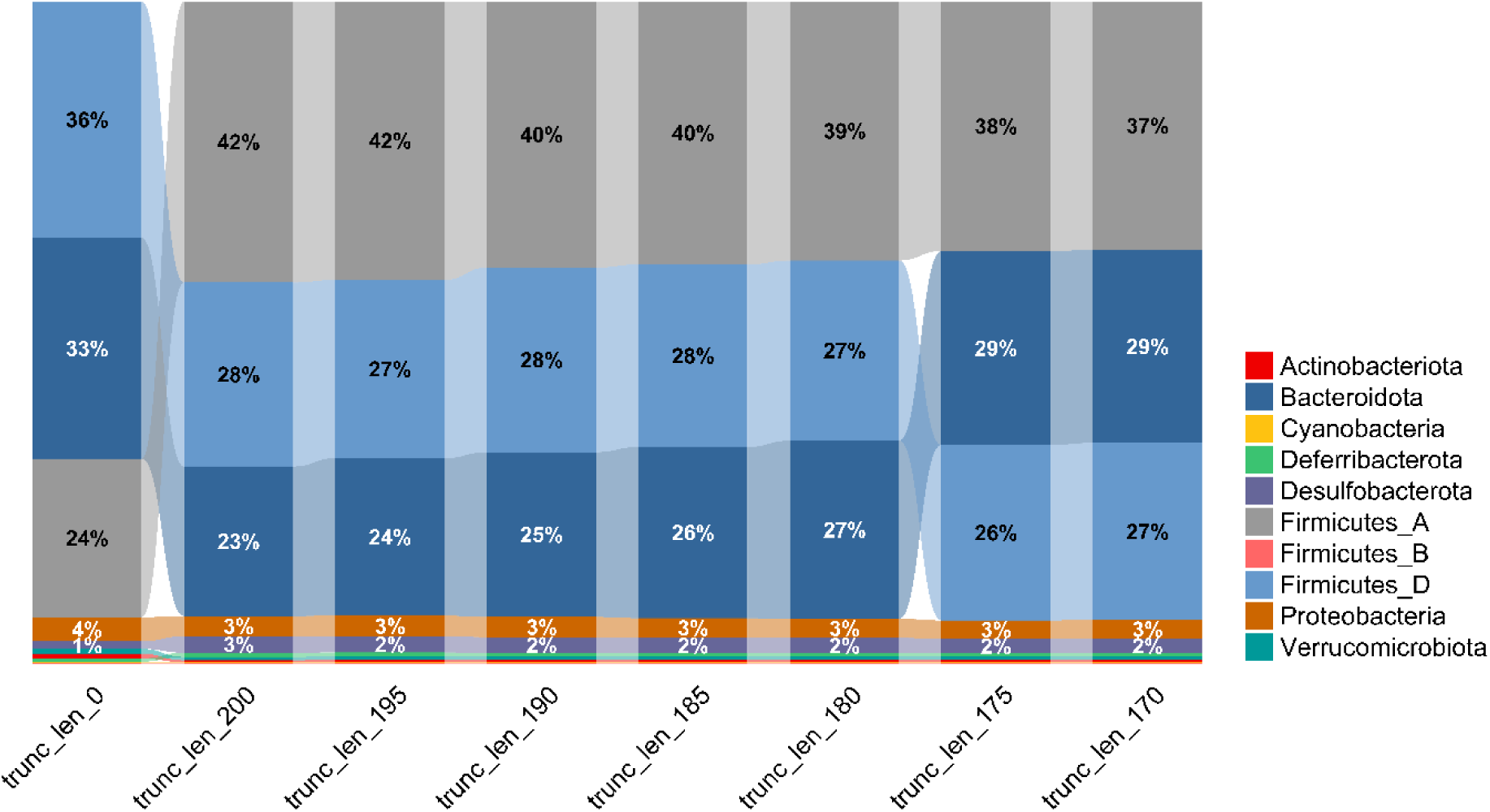
Ribbon chart showing the relative abundance of different truncation lengths at phylum level.

**Fig. 6.**
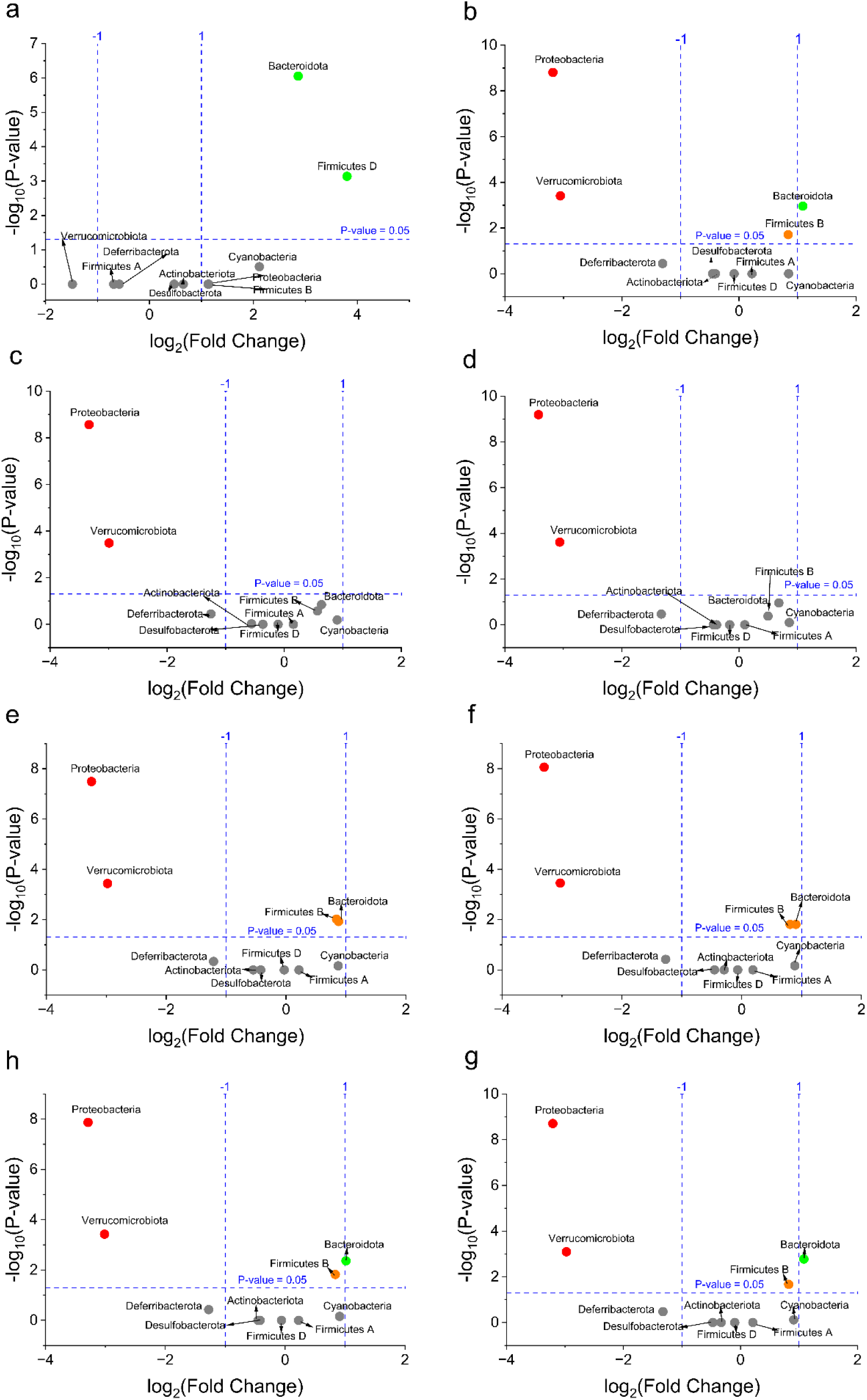
ANCOM volcano plot of differential abundance between group 1 and group 2 data set (**a**) truncation length at 0 (**b**) truncation length at 200 (**c**) truncation length at 195 (**d**) truncation length at 190 (**e**) truncation length at 185 (**f**) truncation length at 180 (**g**) truncation length at 175 (**h**) truncation length at 170.

**Fig. 7.**
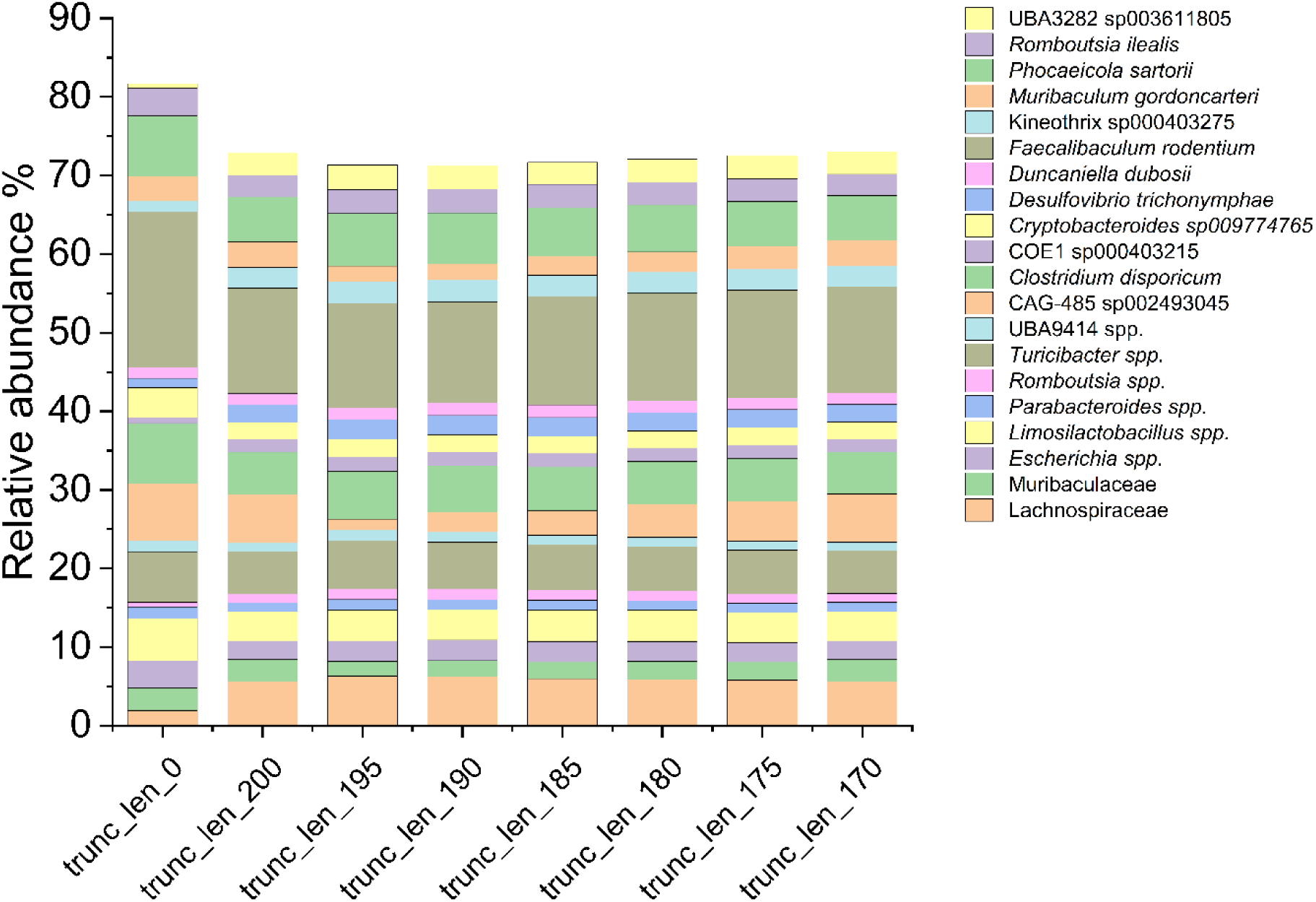
Barplot of relative abundance of different truncation lengths at species level.

## Discussion

The analysis of different truncation lengths in sequencing data is integral to optimizing the accuracy and completeness of microbiome studies. In our research, we systematically evaluated multiple truncation lengths, concluding that trunc_len_175 strikes an ideal balance between maintaining high read recovery rates and preserving microbial diversity. A similar study by Lee et al. [12] demonstrates the trimming condition for DADA2 analysis in QIIME2. In the study, they used the V3-V4 region of human oral microbiota samples and optimized the minimum overlapping length at 16 bps with a Phred quality score of 20. In our study, we have kept the Phred quality score at 18 [13] and the overlapping length was kept as default so that maximum overlap can be achieved and did not compromise with richness of the microbial community. This finding aligns with and expands upon existing literature, demonstrating that careful parameter selection is essential for interpreting microbial community structures effectively.

Previous studies have highlighted the impact of truncation lengths on data quality and community representation. Callahan et al. [2] emphasized that excessive truncation can lead to significant reductions in the number of usable sequences, thereby diminishing the presence of less abundant taxa. Their analysis found that a truncation length of 220 bases optimized read quality while still capturing the richness of the microbial community. Similarly, McMurdie and Holmes [14] found that more stringent truncation adversely affected alpha diversity, as high-throughput sequencing often fails to recover rare taxa that contribute to overall community dynamics, which mirrors our findings where trunc_len_0 produced the most sequences but led to low diversity metrics.

Our results showed that trunc_len_0 yielded the highest overall read counts but was associated with the lowest alpha diversity (Fig. 3a), indicating that while this parameter captured a broad spectrum of reads, it disproportionately favored highly abundant OTUs. This phenomenon is consistent with conclusions drawn from McKnight et al. [15], which noted that untrimmed reads may skew results toward dominant taxa, potentially masking the presence of low-abundance species that could be crucial for ecological insights.

In contrast, trunc_len_200, aimed at improving data quality by filtering out shorter reads, resulted in a significant loss of data. This finding echoes the work of Bokulich et al. [1], who established that overly strict filtering criteria hindered the recovery of diverse microbial populations. Their research illustrated that essential taxa might be omitted if truncation thresholds are set too stringently, emphasizing the importance of balancing quality control with adequate representation of microbial diversity.

Our analysis identified trunc_len_175 as the most effective strategy. This parameter not only maintained a substantial number of filtered and non-chimeric reads (Tab. 1) but also positively impacted the representation of key phyla. For instance, we observed that Bacteroidota became significantly more prominent at this truncation length, which is consistent with findings from Turnbaugh et al. [16]. Their extensive work on the human gut microbiome underscored the importance of appropriately balanced truncation lengths to characterize community structures accurately, especially when working with abundant and rare taxa alike.

Furthermore, the significant shifts in taxa abundance observed at trunc_len_175 align closely with the microbial dynamics reported in studies of similar environments. For example, in a study by Lahti et al. [17] , a balanced approach to truncation directly influenced the clarity and interpretability of the results regarding Bacteroidota and Firmicutes composition in various hosts. The ability to discern these shifts is critical for understanding the ecological roles that different taxa play in health and disease contexts.

## Conclusion

Based on our findings and corroborated by established literature, trunc_len_175 emerges as the best truncation strategy for microbial sequencing studies. This length not only optimizes read recovery but also preserves the richness and evenness of microbial communities, facilitating more nuanced interpretations of ecological dynamics. Future investigations should incorporate these insights and benchmark against established datasets to refine truncation protocols further. By ensuring that methodological choices in high-throughput sequencing are informed by empirical evidence, researchers can enhance the reliability and applicability of microbiome research outcomes.

### Methods Sample data

The sample data set consists of 20 sequences of mice fecal samples selected from our study, 11 samples from the group fed with atherogenic diet (Group1) and nine samples from the group fed with normal chow diet (Group2) (Bioproject no. PRJNA1143515). These samples were sequenced in Illumina MiSeq using 2X250bp paired-end technology. We randomly selected samples of the V4 hypervariable region with accession codes SRS22237203, SRS22237228, SRS22237243, SRS22237244, SRS22237247, SRS22237252, SRS22237255, SRS22237256, SRS22237295, SRS22237296, SRS22237298, SRS22237303, SRS22237304, SRS22237316, SRS22237325, SRS22237326, SRS22237327, SRS22237328, SRS22237329, SRS22237432.

### QIIME2 analysis

QIIME2 [18](Version 2023.7) was used for this study. The raw reads fastq files were imported using the input format “PairedEndFastqManifestPhred33V2” in QIIME2 artifact file .qza. The artifact files were quality filtered, denoised, joined reads, chimera removed, and dereplicated similar sequences into amplicon sequence variants (ASVs) in DADA2 [19]. In the DADA2 analysis, the Phred score was kept at 18 [13], the V4 primer sequences were trimmed, the truncation length was set at 0, 200, 195, 190, 185, 180, 175, and 170 for both forward and reverse reads. The rest of the parameters were set by default. OTU clustering was done using the 16S_full-length Greengenes2 (version 2022.10) database at 99% similarity. Taxonomic classification was performed based on the Naïve Bayes classifier Greengenes2 2022.10 515F/806R V4 region. The classified features were subjected to stringent quality filters. Host DNA including the mitochondria/ chloroplast, features having one or two frequencies, and features and frequencies less than 0.01% were removed during the filtering steps. Microbiome composition (ANCOM) was analysed on the phylum level to determine differential abundance among the groups of various truncation length parameters. The alpha-rarefaction curve was analyzed for the richness matrix with a maximum sequencing depth of 10000, and the rest of the parameters were set by default.

### Alpha diversity and statistical analysis

The feature ASVs table, taxonomy file, and metadata were imported as .csv files in MicrobiomAnalyst 2.0 [20] and alpha diversity index was calculated. Bar graph, box plot, ribbon plot, and volcano plot were prepared using OriginPro, version 2024 (OriginLab Corporation, Northampton, MA, USA.). Alpha rarefaction curve data was imported into R and plotting was done using ggplot2 [21], tidyr [22] and dplyr [23].

## Abbreviation

ASV: Amplicon Sequence variants
DADA: Divisive Amplicon Denoising Algorithm
OTU: Operational Taxonomic Unit
QIIME: Qualitative insights into microbial ecology

## Acknowledgement

This study received funding from the Indian Council of Medical Research (ICMR), New Delhi, India, under its Junior Research Fellowship program, awarded to M.G.S. (JRF No.: 3/1/3/JRF-2019/HRD-070(31035), Project No. GAP0797). The Council of Scientific and Industrial Research (CSIR), New Delhi, and the CSIR-NEIST, Jorhat, Assam, are gratefully acknowledged for providing the necessary facilities, thereby facilitating the smooth progression of the study. The authors also thank the Publication & Intellectual Property Rights Committee, CSIR-NEIST, Jorhat for approving the manuscript for publication (Manuscript No: CSIR-NEIST/PUB/2025/026, dated 11-02-2025).

## Author contribution

W.R. developed the detailed protocols and oversaw all stages of the study, including critical inputs and the manuscript drafting process. M.G.S. conceptualized the study, performed data analysis, generated the empirical data, and wrote the manuscript. All authors reviewed and edited the final draft of the manuscript and collectively approved the decision to submit the manuscript for publication consideration.

## Availability of data and materials

Raw data are available in NCBI SRA under Bioproject No. PRJNA1143515

## Ethics approval and consent to participate

Not applicable

## Consent for publication

Not applicable

